# Quantitative Proteomic Analysis of Prostate Tissue Specimens Identifies Deregulated Protein Complexes in Primary Prostate Cancer

**DOI:** 10.1101/447375

**Authors:** Bo Zhou, Yiwu Yan, Yang Wang, Sungyong You, Michael R. Freeman, Wei Yang

## Abstract

Prostate cancer (PCa) is the most frequently diagnosed non-skin cancer and a leading cause of mortality among males in developed countries. However, our understanding of the global changes of protein complexes within PCa tissue specimens remains very limited, although it has been well recognized that protein complexes carry out essentially all major processes in living organisms and that their deregulation drives the pathogenesis and progression of various diseases. By coupling tandem mass tagging-synchronous precursor selection-mass spectrometry/mass spectrometry/mass spectrometry (TMT-SPSMS3) with differential expression and co-regulation analyses, the present study compared the differences between protein complexes in normal prostate, low-grade PCa, and high-grade PCa tissue specimens. Globally, a large downregulated putative protein-protein interaction (PPI) network was detected in both low-grade and high-grade PCa, yet a large upregulated putative PPI network was only detected in high-grade but not low-grade PCa, compared with normal controls. To identify specific protein complexes that are deregulated in PCa, quantified proteins were mapped to protein complexes in CORUM, a collection of experimentally verified mammalian protein complexes. Differential expression analysis suggested that mitochondrial ribosomes and the fibrillin-associated protein complex were significantly overexpressed, whereas the ITGA6-ITGB4-Laminin10/12 and the P2X7 receptor signaling complexes were significantly downregulated, in PCa compared with normal prostate. Moreover, differential co-regulation analysis indicated that the assembly levels of some nuclear protein complexes involved in RNA synthesis and processing were significantly increased in low-grade PCa, and those of mitochondrial complex I and its subcomplexes were significantly increased in high-grade PCa, compared with normal prostate. In summary, the study represents the first global and quantitative comparison of protein complexes in prostate tissue specimens. It is expected to enhance our understanding of the molecular mechanisms underlying PCa development and progression in human patients, as well as lead to the discovery of novel biomarkers and therapeutic targets for precision management of PCa.

## Background

Prostate cancer (PCa) is the most frequently diagnosed non-skin cancer and a leading cause of cancer death among males in developed countries [1]. In the United States alone, it was estimated that 164, 690 men will be diagnosed with PCa and that 29, 430 will die of this disease in 2018 [2]. Largely owing to the developments and advances in next-generation sequencing technologies, the past few years have witnessed a striking growth of genomic and transcriptomic profiles of clinical PCa specimens [3]. These large-scale efforts not only enhanced our understanding of the molecular underpinnings of PCa pathogenesis and progression, but also facilitated the identification of novel PCa biomarkers and therapeutic targets [4, 5]. Nonetheless, despite the significant progress, genomic and transcriptomic profiling studies have inherent limitations — they only indirectly and often inconclusively measure the properties of proteins, which are the major functional molecules and actual executors of biological functions in living organisms. In fact, recent studies have suggested that aberrations at the gene copy number, DNA methylation, and RNA expression levels often do not reliably predict changes at the protein expression level [6, 7].

In contrast to genomic and transcriptomic technologies, mass spectrometry (MS)-based proteomic technologies enable comprehensive and direct analysis of proteins, and have thus been widely used in the proteomic profiling of clinical specimens, such as biofluid and tissue samples [8]. Compared with biofluid specimens such as blood and urine, tissue specimens allow more accurate sampling of proteomic changes in tumor cells and microenvironment, but they are more difficult to obtain. According to a recent survey, only about 40 proteomic studies were performed on human PCa tissue specimens in the past decade [9]. Moreover, most of these studies were conducted using the two-dimensional electrophoresis (2DE) matrix-assisted laser desorption ionization mass spectrometry (MALDI-MS) technology, which rarely provides adequate proteomic coverage and is only semi-quantitative. To date, comprehensive and quantitative proteomic studies of PCa tissue specimens have remained scarce [6, 10–14]. Furthermore, none of these studies investigated the global changes of multiprotein complexes along PCa development and progression. Notably, protein complexes act as highly specialized molecular machines and carry out essentially all major processes in a cell, such as gene transcription and splicing as well as protein synthesis and degradation [15, 16]. The abnormal expression and/or activation of certain protein complexes may lead to the pathogenesis and progression of many diseases [17]. Hence, the identification of deregulated protein complexes in clinical tissue specimens offers a great potential of revealing novel molecular mechanisms and discovering new biomarkers and therapeutic targets for various human diseases including PCa.

Currently, a variety of proteomic technologies are available for large-scale protein quantification [18]. Among these, tandem mass tagging (TMT) offers high multiplexing capability, allowing quantitative comparison of up to 11 samples simultaneously [19, 20]. Previously, TMT suffered from the issue of precursor ion interference, which results in ratio compression and thus an underestimation of expression differences [21]. With the recent development of the synchronous precursor selection (SPS)-MS3 technique, the ratio compression issue is largely eliminated [22]. As such, the TMT-SPS-MS3 combination enables highly multiplexed and accurate quantification of proteomes. Indeed, a recent study showed that TMT-SPS-MS3 enables more accurate protein quantification than label-free quantification, especially for modest (<2-fold) changes [23]. Moreover, when coupled with protein co-regulation analysis, TMT-SPS-MS3 analysis permits systems-wide analysis of protein-protein associations with high accuracy [24].

In the present study, the TMT-SPS-MS3 approach was integrated with differential expression and co-regulation analyses to investigate the global changes of protein complexes, in terms of abundance and protein-protein associations, in 27 optimal cutting temperature (OCT) compound-embedded and cryopreserved clinical tissue specimens of primary PCa (*i.e.*, 9 normal prostate, 9 low-grade/low-risk PCa, and 9 high-grade/high-risk PCa). Notably, recent studies have shown that, when properly handled and processed, OCT samples provide better protein recovery and MS identification than formalin-fixed and paraffin-embedded (FFPE) specimens [25, 26]. After stringent statistical analysis, the study revealed that certain protein complexes were significantly deregulated in low-grade and/or high-grade PCa, compared with normal prostate. Further exploitation of the deregulated protein complexes may shed new light on the molecular basis of PCa development and progression *in vivo*, as well as provide novel biomarkers and therapeutic targets for better management of this leading male cancer.

## Methods

### Prostate tissue specimens

All PCa and PCa-adjacent normal tissue samples were collected from radical prostatectomy at Cedars-Sinai Medical Center, during the period of 2010 to 2014 (Table S1). The PCa samples are either of low-grade (Gleason score of 6) or high-grade (Gleason score of 8 or 9) prostate adenocarcinoma. All specimens were OCT embedded and stored at −80°C prior to proteomic analysis.

### Protein extraction, digestion and TMT labeling

OCT was removed essentially as described [27]. Briefly, about 20 mm^3^ tissue was cut into small pieces and transferred to 1.5 mL Eppendorf tubes. Tissue pieces were gently washed with 1 mL ice-cold 70% ethanol for twice, ice-cold water for once, and ice-cold 100 mM Tris-HCl, pH 7.4 for twice. To lyse tissue, 100 μL lysis buffer (80 mM Tris-HCl, 4% SDS, 100 mM DTT, pH7.4) was added into each tube, and the tissue pieces were grinded with disposable pestles using a cordless pestle motor (VWR, Radnor, PA). The lysates were thoroughly sonicated in a water-bath sonicator (Elma S180H) to reduce viscosity, incubated at 95°C for 5 min, and centrifuged at 16,000×g for 10 min. Protein concentration was determined using the Pierce 660 nm protein assay (Thermo Scientific) according to the manufacture’s instruction. To generate an internal proteomic standard, 20 μg protein from each of the 27 samples was mixed. Because a 10-plex TMT reagent set can only accommodate up to 10 samples, the 27 tissue samples and three internal standard (pooled) samples were divided into three sets. Each set contains one internal standard, three normal prostate, three low-grade PCa, and three high-grade PCa samples. From each sample, 60 μg proteins were alkylated with iodoacetamide and digested with trypsin using the filter-aided sample preparation (FASP) method [27]. Tryptic peptides were labeled with 10-plex TMT reagents in parallel, essentially as we previously described [28, 29]. To ensure that the internal standards for the three sets are identical, the three TMT126-labeled internal standard samples were mixed into one single sample. Subsequently, for each TMT10plex set, an equal amount of tryptic peptides (derived from about 20 μg proteins) with differential TMT labeling was merged into one sample, desalted using C_18_ spin columns(Thermo Scientific), and concentrated in a SpeedVac (Thermo Scientific).

### Peptide fractionation

Peptide fractionation was performed using high-pH reversed-phase liquid chromatography (LC) [30]. Each TMT10plex-labeled peptide mixture sample was redissolved with 45 μL 10 mM ammonium formate, pH 10. Twenty microliters of peptide solution were injected and separated on a 20-cm Hypersil GOLD C_18_ column (1.9 μm particle size, 2.1 mm inner diameter, 175 Å pore size) heated to 35°C on an Ultimate 3000 XRS system (Thermo Scientific), with a flow rate of 0.5 mL/min. Mobile phase A and B consisted of 10 mM ammonium formate in water (pH 10) and 10 mM ammonium formate in 95% acetonitrile (pH 10), respectively. The 13-min LC gradient was 0% B over 3 min, 0-28% B over 7 min, 28-90% B over 1 min, 90% B over 1 min, and 90-0% B over 1 min. For each TMT10plex set, a total of 72 fractions were collected after 3.5 min, with a collection rate of one fraction per 6 sec. The 72 fractions were then concatenated into 24 fractions by combining fractions 1, 25, 49; 2, 26, 50; and so on. The fractions were concentrated in a SpeedVac and stored at −80°C until LC-SPS-MS3 analysis.

### LC-SPS-MS3 analysis

LC-SPS-MS3 analysis was conducted on an EASY nLC 1200 connected to an Orbitrap Fusion Lumos mass spectrometer (Thermo Scientific). Each fraction of peptides was redissolved with 25 μL 0.2% formic acid, 2% acetonitrile. Ten microliters of peptide solution were loaded onto a 2-cm trap column (PepMap 100 C_18_, 75 μm inner diameter, 3 μm particles, 100 Å pore size) and separated by a 50-cm EASY-Spray column (PepMap RSLC C_18_, 75 μm inner diameter, 2 μm particles, 100 Å pore size) heated to 55°C, at a flow rate of 250 nL/min. Mobile phases A and B consisted of 0.1% formic acid in water and 0.1% formic acid in 80% acetonitrile, respectively. The 3-h LC gradient was 3-25% B over 140 min, 25-50% B over 25 min, 50-100% B over 5 min, and 100% B over 10 min. SPS-MS3 analysis was conducted essentially as described [23]. The parameter settings for FTMS1 include orbitrap resolution (120, 000), scan range (350-1400), AGC (5E5), maximum injection time (100 ms), RF lens (30%), data type (centroid), charge state (2-5), dynamic exclusion for 60 s using a mass tolerance of 7 ppm, and internal calibration using m/z 371.10123; for ITMS2 include mass range (400-1400), number of dependent scans (10), isolation window (0.4 m/z), activation type (rapid CID), collision energy (35%), maximum injection time (120 ms), AGC (2E4), and data type (centroid); for MS3 include mass range (400-1400), precursor ion exclusion (low m/z 50, high m/z 5), isolation window (m/z 0.7), MS2 isolation window (m/z 2), number of notches (10), HCD collision energy (55%), orbitrap resolution (50,000), maximum injection time (150 ms), AGC (2.5E5), and data type (centroid).

### Protein Identification and Quantification

The acquired LC-SPS-MS3 files were analyzed by MaxQuant (v1.6.0.16) [31], using the Andromeda algorithm [32] to search against the human Uniprot protein sequence database (released on 03/30/2018, containing 20,937 canonical sequences and 72,379 additional sequences) combined with the common contaminant protein sequences (244 sequences). The quantification type was Reporter ion MS3, the isobaric labels were TMT10plex, and the reporter mass tolerance was 0.003 Da. Modifications included carbamidomethylation of cysteines as fixed modification as well as acetylation of protein N-terminus, deamidation of asparagines and glutamines, and oxidation of methionines and prolines as variable modifications. Tryspin/P was used for digestion and up to two miscleavages were allowed. The match-between-runs function was enabled, using 0.7 min of match time window and 20 min of alignment time window. The mass tolerance was 20 ppm for first search peptide tolerance and 4.5 ppm for main search peptide tolerance, and 0.5 Da for MS/MS match tolerance. A false discovery rate (FDR) of 1% was applied to filter peptide-spectrum matches (PSMs), peptides, and protein groups. The mass spectrometry proteomics data have been deposited to the ProteomeXchange Consortium (http://proteomexchange.org) via the PRIDE partner repository [33] with database identifier PXD010744.

### Identification of differentially expressed proteins (DEPs)

Statistical analysis was performed with Perseus (v1.5.5.3) [34]. Proteins identified from the reverse sequence database or based on a single modified peptide, as well as non-human contaminant proteins identified from the contaminant sequence database, were filtered out. Subsequently, only proteins quantified across all the analyzed samples were selected for statistical analysis. Protein ratios against the internal standard (TMT126 channel) were computed and then log_2_-transformed. For each sample, the log_2_-transformed ratios were normalized against the Tukey’s bi-weight mean, which calculates a robust average that is unaffected by outliers, with the assumption that most identified proteins are not significantly differentially expressed across the samples. After a quality control analysis using SuperHirn [35], one outlier sample was detected from each group and they were removed. For the comparison between each group (n = 8 after the removal of outlier samples), Student’s *t*-test (two-tailed) was used. To correct the *p* values for multiple testing, the Storey method was applied [36]. DEPs were identified using *q* values < 0.05 and the empirical cutoff of log_2_-transformed fold changes of > 0.5 in absolute value. For gene ontology enrichment, the Database for Annotation, Visualization and Integrated Discovery (DAVID, v6.8) analysis was performed [37]. To generate putative protein-protein interaction (PPI) networks from DEPs, the Ingenuity Pathway Analysis (Ingenuity) was performed with high stringency — only direct PPIs with experimental evidence were used.

### Identification of differentially expressed protein complexes

To identify specific protein complexes that are differentially expressed, the CORUM annotation for each protein was added to the data matrix in Perseus (v1.5.5.3), followed by the extraction of CORUM complexes containing at least five quantified proteins using R statistical software (R Development Core Team; https://www.r-project.org/) (v3.5.0). The mean log2-ratio of all proteins in a CORUM complex was calculated for each sample, and then compared across the three groups (*i.e.*, normal, low-grade, and high-grade) by Student’s *t*-test (two-tailed). The CORUM complexes with *p* values of <0.05 and the mean difference of > 0.3 in absolute value were accepted as differentially expressed complexes. Here, the cutoff for the mean difference was set as 0.3 because it corresponds to p < 0.05, based on the normal distribution of all mean differences of low-grade PCa and high-grade PCa versus normal controls (s.d. = 0.142).

### Identification of differentially regulated protein complexes

Differential co-regulation analysis provides a level of information about protein-protein associations, which is not possible to obtain using the widely used differential expression analysis [24, 38]. Differential co-regulation analysis was conducted using R (v3.5.0) to analyze the pairwise correlation of proteins within each CORUM complex. The Spearman’s method was used to assess correlation of proteins within each complex. Subsequently, the Fisher z-transformation was performed to stabilize the variance of sample correlation coefficients in each condition, as described in [39, 40]. To avoid obtaining infinite z scores, all Spearman’s Rho values of 0.99 through 1 were replaced by 0.99 and those of −1 through −0.99 were replaced by −0.99. For each CORUM complex, to determine whether the difference of mean z scores between two conditions (*e.g.*, normal prostate vs low-grade PCa) is statistically significant, the following steps were performed: 1) 8 out of the 24 samples were randomly sampled twice and used as condition A and condition B, respectively; 2) for each condition, Spearman’s Rho values and z scores were computed as mentioned above; 3) the mean z score difference between conditions A and B was calculated; 4) the steps 1 to 3 were repeated for 10,000 times, and null hypothesis distribution of the mean z score differences was generated; and 5) the significance of each observed mean z score difference was computed using the null hypothesis distribution. CORUM protein complexes with *p* values of < 0.05 and mean z score differences of > 0.5 in absolute value were accepted as differentially regulated protein complexes. As the standard deviation of all mean z score differences of low-grade PCa and high-grade PCa versus normal was 0.324, the cutoff for the mean z score difference of 0.5 corresponds to *p* < 0.1.

## Results

### TMT-SPS-MS3 analysis for quantitative profiling of prostate tissue specimens

The TMT-SPS-MS3 method was applied to quantitatively compare the proteomes across the 27 OCT-embedded prostate tissue samples, followed by differential expression and co-regulation analyses to identify *in vivo* deregulated protein complexes (Fig. 1). The specimens were divided into three risk groups according to the prostatectomy Gleason scores, including normal prostate (abbreviated as N, n=9), low-grade PCa (LG, n=9), and high-grade PCa (HG, n=9). Of note, the Gleason score (on a scale of 6 to 10) is one of the most commonly used systems for evaluating the aggressiveness of primary PCa, and a higher Gleason score is generally associated with a worse prognosis [41, 42].

**Figure 1.**
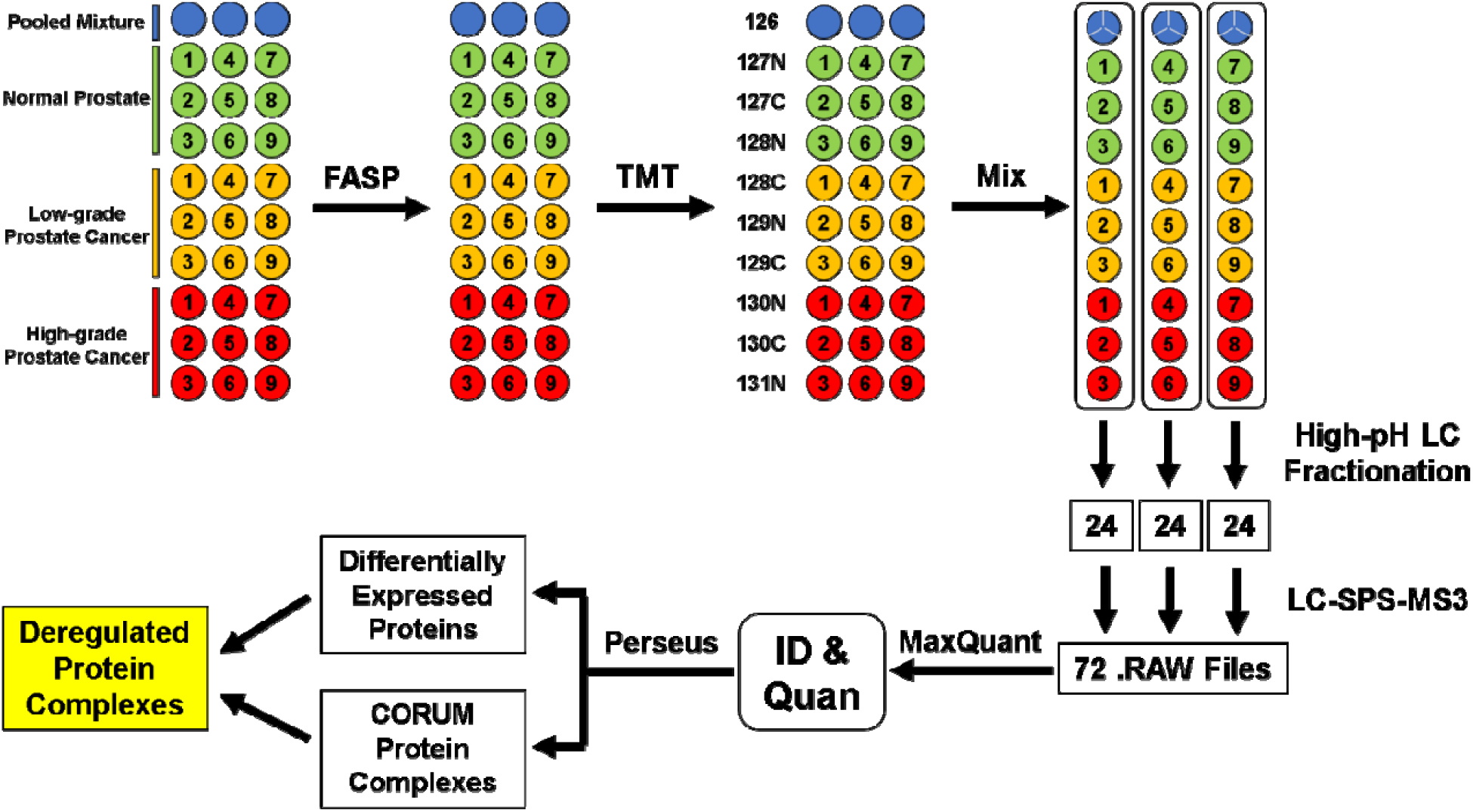
Workflow for quantitative proteomic comparison of three groups of prostate tissue ue specimens (*i.e.*, normal control, low-grade PCa, and high-grade PCa) using TMT-SPS-MS3. A totall of 30 OCT-embedded prostate tissue samples were digested in parallel into tryptic peptides by FASP, followed by chemical labeling with three sets of TMT10plex reagents. The three TMT126-labeled pooleded mixture samples (shown as blue circles) were mixed and then equally divided into three portions (shown n as blue circles divided into thirds). Differentially TMT-labeled peptide samples were mixed, and then en each TMT10plex mixture sample was fractionated into 72 fractions by high-pH RPLC and concatenated ed into 24 fractions. Each fraction of peptides was analyzed by LC-SPS-MS3. The acquired 72 RAW files s were analyzed by MaxQuant to identify and quantify proteins. Proteins quantified across all the 30 samples were used for differential expression and co-regulation analyses to identify *in vivo* deregulated protein complexes.

### Identification and analysis of DEPs

After TMT-SPS-MS3 analysis, database searching, and protein identification filtering, a total of 5,562 protein groups were identified with an FDR of ≤1%. Among these, 5,297, 4, 662, and 3,642 protein groups were quantified in at least one, two, and three TMT sets, respectively (Fig. S1 and Table S2). The 3,642 protein groups that were quantified across all the 27 samples were used for the following statistical and bioinformatic analyses.

Quantified profiles were examined degree of variation between samples for quality assessment using superHirn [35], and samples with high degree of between-sample variation were removed prior to further statistical analysis (Fig. S2). For the remaining 24 samples (n=8 for each group), after Student’s t test (two-tailed) and multiple comparison correction, the cutoffs of *q* <0.05 and log2-ratios of > 0.5 in absolute value were applied to identify DEPs (Table S3). A total of 197 DEPs, including 143 downregulated and 54 upregulated proteins, were identified in LG samples in comparison to the N samples (Fig. 2A and Table S4). Of these, MAM domain-containing protein 2 (MAMDC2) and pyrroline5-carboxylate reductase 1 (PYCR1) were the most dramatically downregulated and upregulated proteins, respectively (Fig. 2A, lower panel). In HG samples (versus N samples), a total of 309 DEPs (215 downregulated and 94 upregulated) were identified, among which glutathione S-transferase mu 1 (GSTM1) and spondin-2 (SPON2) were the most remarkably downregulated and upregulated proteins, respectively (Fig. 3B and Table S4). In comparison, the proteomic difference between LG and HG samples was small — only 33 DEPs (29 downregulated and 4 upregulated in HG, compared with LG) were identified, of which ADP-ribosyl cyclase/cyclic ADP-ribose hydrolase 1 (CD38) and decaprenyl diphosphate synthase subunit 2 (PDSS2) were the most substantially downregulated and upregulated proteins, respectively (Fig. 3C and Table S5).

**Figure 2.**
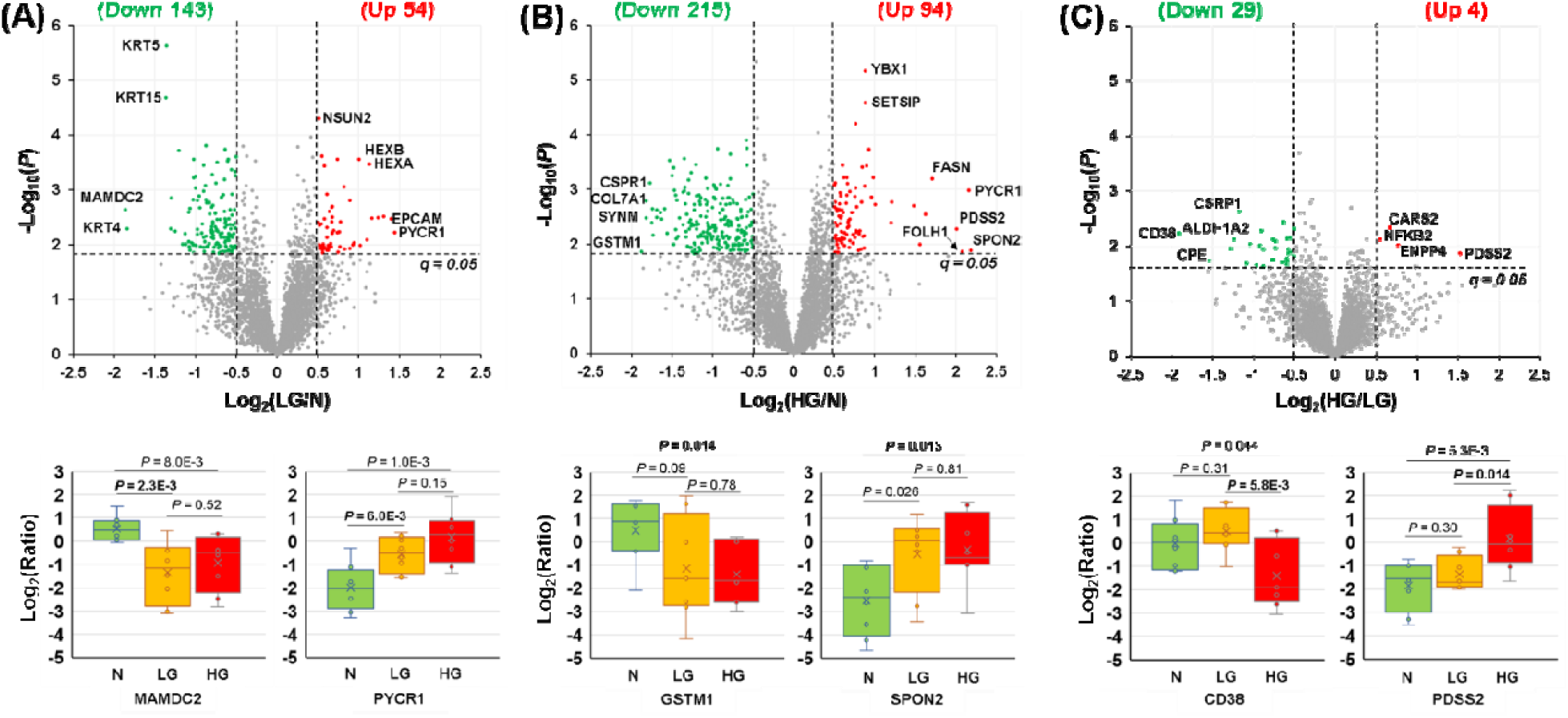
Identification of DEPs between (A) N vs LG, (B) N vs HG, and (C) LG vs HG. The upperer panel shows the volcano plots of all the 3,642 quantified protein groups, and the lower panel shows the he boxplots for the most remarkably changed proteins in each comparison. Here, the abbreviations DEP,N,N,LG, and HG stand for differentially expressed proteins, normal, low-grade, and high-grade, respectively.

**Figure 3.**
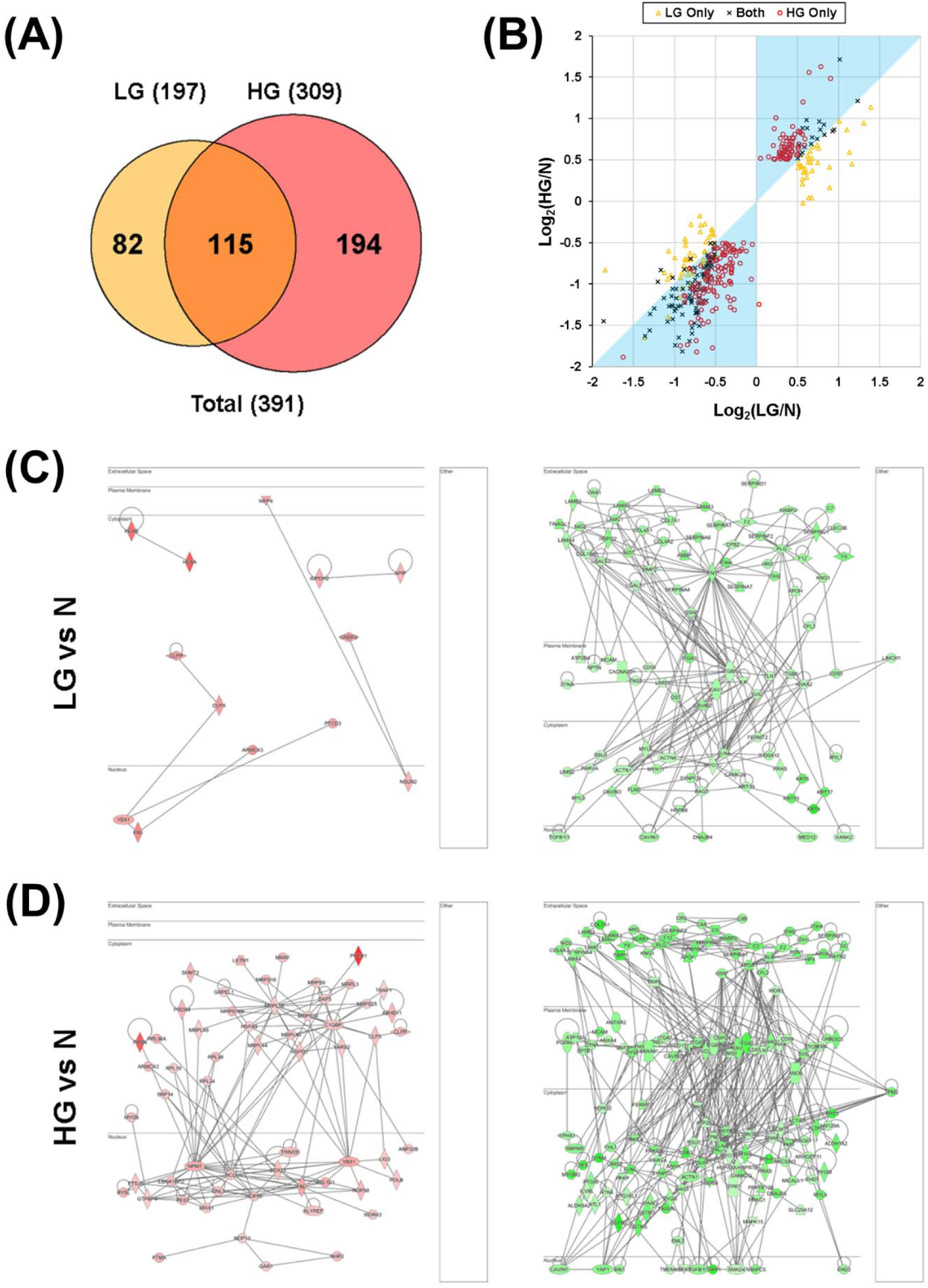
Comparison of protein groups differentially expressed in LG and HG PCa, compared with N samples. **(A)** Venn diagram of protein groups differentially expressed in LG and HG PCa, compared with N samples. A total of 82 DEPs were LG only, 194 were HG only, and 115 were shared by both LG and HG. **(B)** Scatter plot for the comparison of log_2_(LG/N) and log_2_(HG/N) ratios for the 82 LG-only, 115 shared, and 194 HG-only DEPs. The cyan shade covers the area where the absolute values of log_2_(LG/N) are less than those of log_2_(HG/N), *i.e.*, the changes in the LG group are less remarkable than those in the HG group. **(C)** Putative networks of direct PPIs for proteins significantly upregulated (left) or downregulated (right) in LG PCa, compared with N samples. The four subcellular localization layers are extracellular space, plasma membrane, cytoplasm, and nucleus (from top to bottom). **(D)** Putative networks of direct PPIs for proteins significantly upregulated (left) or downregulated (right) in HG PCa, compared with N samples.

A comparison of the 197 DEPs in the LG (vs N) group and the 309 DEPs in the HG (vs N) group suggested that 115 DEPs are shared by the two groups, whereas 82 and 194 are unique to the LG and HG groups, respectively (Fig. 3A and Table S4). Interestingly, most of the 194 DEPs unique to the HG group were also differentially expressed in the LG group, albeit to a lesser degree (with log_2_-ratios of −0.5 to 0.5) (Fig. 3B, red circles). Similarly, most of the 115 shared DEPs were less changed in the LG group than in the HG group (Fig. 3B, black “x”s). Collectively, the findings suggest that, compared with normal prostate, most protein expression level changes that are statistically significant in high-grade PCa were already present in low-grade PCa samples, although the extent is less pronounced. In contrast, most of the 82 DEPs unique to the LG group were less markedly changed in the HG group (Fig. 3B, orange triangles). Intriguingly, DAVID analysis of the 82 LG-only DEPs revealed a highly significant over-representation of extracellular exosomes (53 proteins, *p* = 2E-24) (Table S6), suggesting the possibility of differential exosome biogenesis and/or shedding in the two groups.

To determine whether the DEPs may interact with each other and form protein complexes, IPA was applied to reconstruct networks of proteins with direct PPI evidence that was experimentally obtained. In the LG group (vs N), only 13 out of the 54 (24%) upregulated DEPs form three small PPI networks, but 92 out of the 143 (64%) downregulated DEPs form a large PPI network (Fig. 3C). In comparison, in the HG group (vs N), 58 out of the 94 (62%) upregulated DEPs form a large upregulated PPI network, and 157 out of the 215 (73%) downregulated DEPs form a large downregulated PPI network (Fig. 3D). Notably, the upregulated protein subnetworks are almost exclusively localized in cytoplasm and nucleus, whereas the downregulated protein subnetworks are mainly localized in extracellular space, plasma membrane, and cytoplasm.

### Identification of differentially expressed CORUM protein complexes

The CORUM database is a manually curated repository of experimentally characterized protein complexes from mammalian organisms, especially human [43]. To identify specific protein complexes that are deregulated in primary PCa, the 3,642 quantified protein groups were mapped to the CORUM database. A total of 179 protein complexes were found to contain at least five proteins in each complex, and they were selected for further analysis (Table S7). To identify differentially expressed CORUM protein complexes, the log_2_ -ratios of proteins in each complex were averaged for each tissue sample, followed by Student’s *t* test for the comparison of LG vs N groups and of HG vs N groups. After applying the cutoffs of *p* < 0.05 and the log_2_-ratios of > 0.3 in absolute value, six complexes (four up and two down) were found to be differentially expressed in the LG group (vs N), and 11 complexes (eight up and three down) were differentially expressed in the HG group (vs N) (Fig. 5A, Fig. 5B, and Table S8). Of note, the six complexes found in the LG group are a part of the 11 complexes found in the HG group (Fig. 5B).

**Figure 5.**
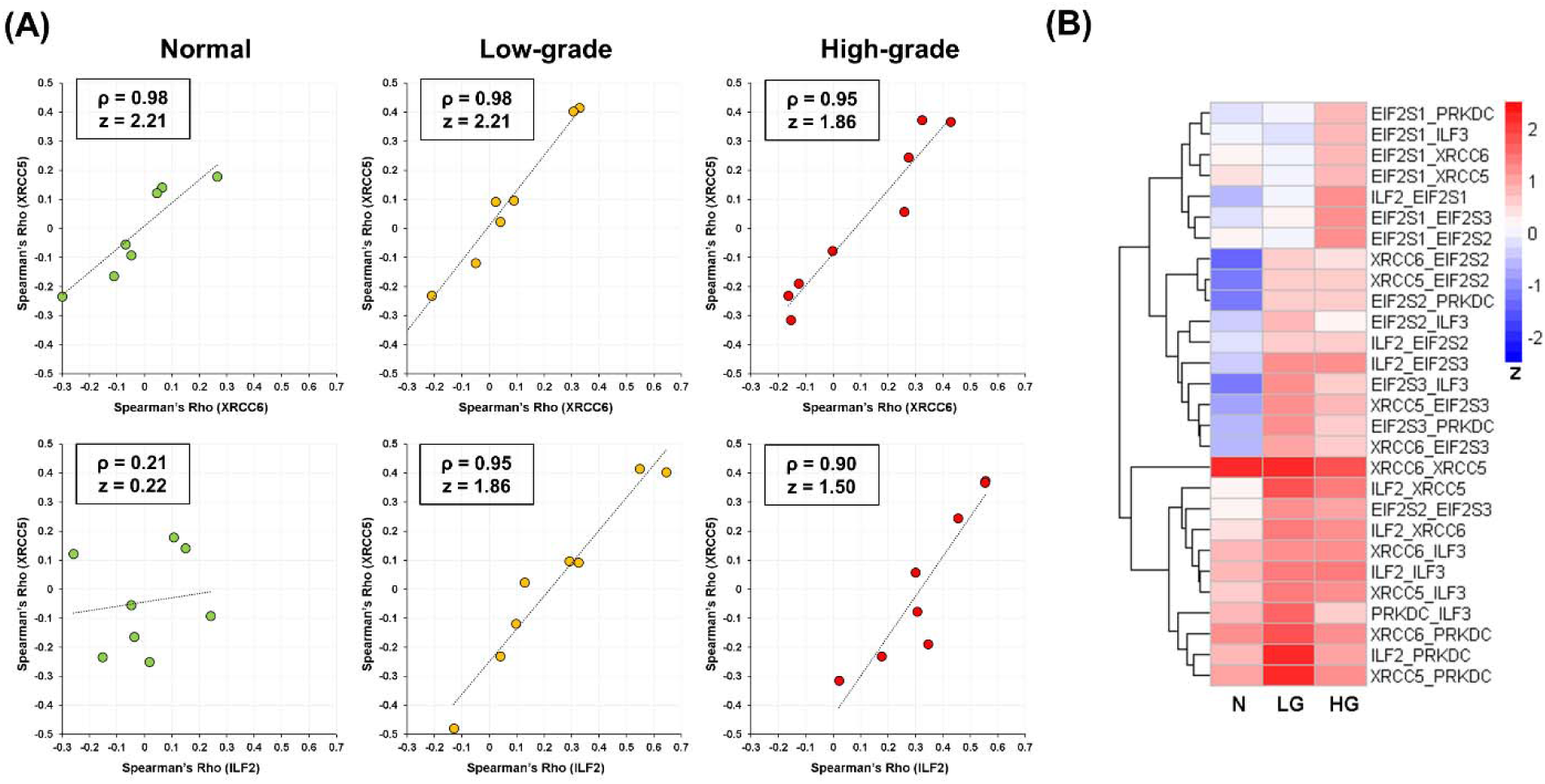
representative example showing the differences of specific protein pairs in the DNA-PK-Ku-eIF2-NF90-NF45 protein complex across the three sample groups. **(A)** Scatter plots showing the Spearman’s Rho of XRCC6 plotted against that of XRCC5 (upper panel) as well as the Spearman’s Rho of ILF2 plotted against that of XRCC5 (lower panel) in the N, LG, and HG groups (from left to right). **(B)** Heatmap of z scores of all protein pairs (excluding self-pairs) within the DNA-PK-Ku-eIF2NF90-NF45 protein complex.

The most upregulated protein complexes are mitochondrial ribosomes, including the entire 55S ribosome as well as the 39S large subunit and the 28S small subunit (Fig. 5B-5D). Other overexpressed protein complexes include two protein complexes involved in ribosome biogenesis (*i.e.*, the fibrillarin-associated protein complex and the Nop56p-associated pre-rRNA complex), a complex involved in RNA splicing (the SRm160/300 splicing coactivator complex), and the cytoplasmic ribosomes (the entire ribosome and the 60S ribosomal subunit). The most downmodulated protein complex is the ITGA6-ITGB4-Laminin10/12 complex, an integrin-laminin complex important for cell-matrix adhesion (Fig. 5B-5D). The other downregulated complexes include the Polycystin-1 multiprotein complex, which plays a key role in focal adhesion, and the P2X7 receptor signaling complex.

### Identification of differentially regulated CORUM protein complexes

Protein-protein associations are essential for the assembly of functional protein complexes. Recent studies provided compelling evidence that co-regulation analysis of protein pairs permits the analysis of protein-protein associations with high accuracy [24, 38]. To identify differentially associated/assembled protein complexes *in vivo*, differential co-regulation analysis was performed for the aforementioned 179 CORUM complexes.

Firstly, for each CORUM complex, the Spearman correlation of each protein pair was computed and then converted into a z score to stabilize the variance [39, 40]. For instance, the DNA-PK-Ku-eIF2-NF90-NF45 complex, which plays a critical role in DNA double-strand break repair, is formed after the Ku heterodimer binds to a suitable DNA end [44, 45]. As expected, the XRCC5 and XRCC6 proteins — subunits of the stable Ku heterodimer [45] — have high Spearman’s Rho values and z scores in all the N, LG, and HG groups (Fig. 5A, upper panel). In comparison, the ILF2 and XRCC5 protein pair has low Spearman’s Rho and z score in the N group but has high values in the LG and HG groups (Fig. 5A, lower panel). This suggests increased protein-protein association between ILF2 and XRCC5 in the LG and HG groups, compared with the N group. A heatmap visualization of the z scores of all protein pairs within the complex showed that most protein pairs have higher z scores in the LG and HG groups than in the N group (Fig. 5B). This indicates that more DNA-PK-Ku-eIF2-NF90-NF45 complexes may be assembled in PCa cells than in normal prostate cells, probably in response to higher DNA damage in cancer cells.

Secondly, to statistically compare the assembly levels of each CORUM protein complex across the three groups, the z scores for all protein pairs within a complex were averaged and the mean z scores were used to estimate the assembly levels of the protein complexes. As shown in Fig. 6A, the mean z scores of all the 179 protein complexes in the N group roughly follow a normal distribution. In comparison, both the LG and HG groups have shoulder peaks on the right side, suggesting that certain protein complexes may have stronger protein-protein associations (*i.e.*, higher assembly levels) in the LG and/or HG groups than in the N group. Furthermore, a heatmap analysis showed that the protein complexes changed in the LG group are mostly different from those changed in the HG group, in comparison to the N group (Fig. 6B).

**Figure 6.**
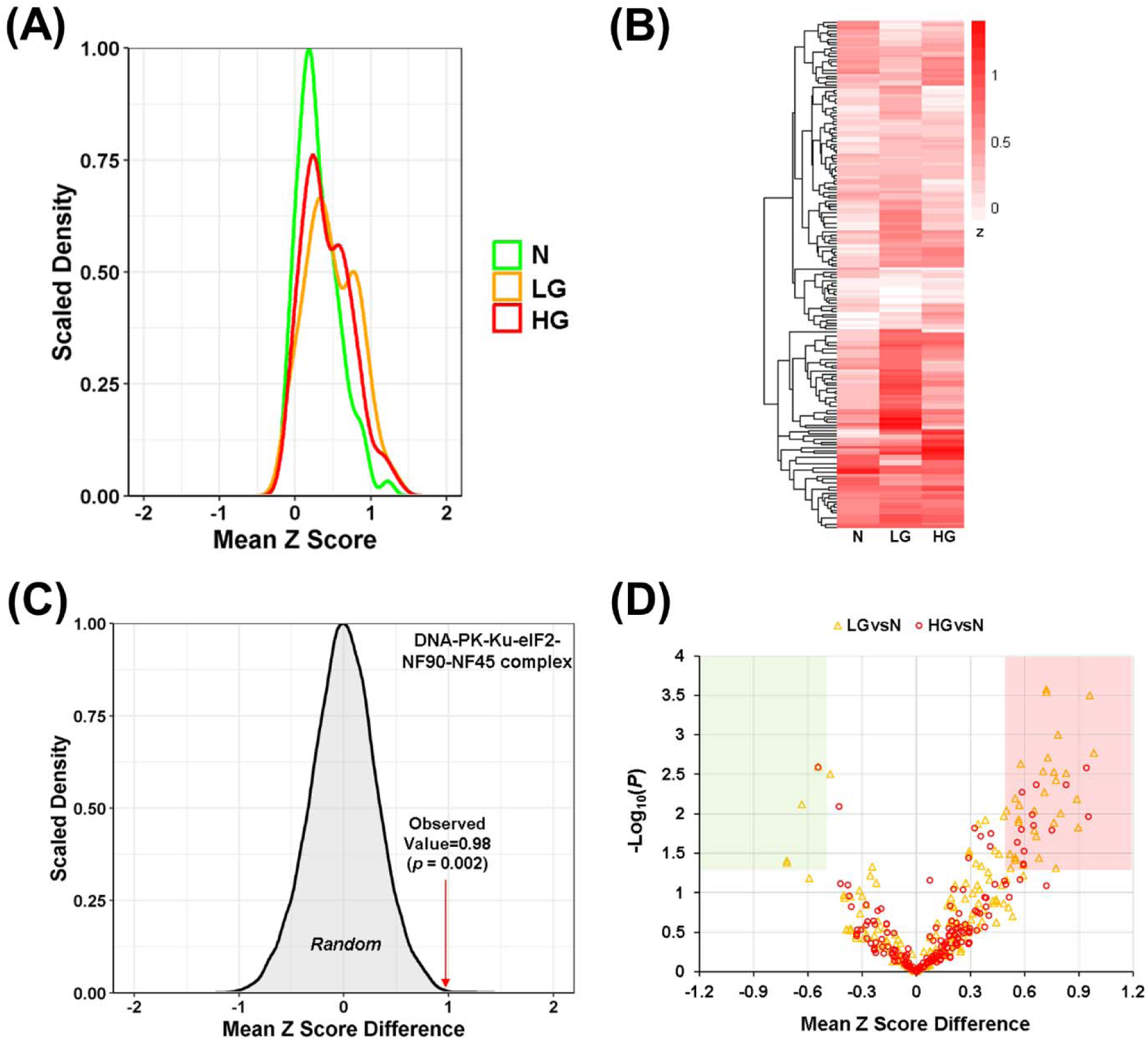
Identification of differentially associated CORUM protein complexes. **(A)** Density plot of the mean z scores for the 179 CORUM protein complexes in N, LG, and HG samples. **(B)** Heatmap of the mean z scores for the 179 CORUM protein complexes in N, LG, and HG specimens. **(C)** A representative example for the determination of *p* value corresponding to an observed mean z score difference. The peak shows the distribution of 10,000 mean z score differences between two sets of eight randomly selected samples. **(D)** Volcano plot showing mean z score differences plotted against negative log_10_-transformed *p* values for the 179 CORUM protein complexes.

Thirdly, for each CORUM protein complex, the distribution of 10,000 random mean z score differences was plotted, and the *p* value corresponding to an observed difference of mean z score between two groups was calculated. For example, for the DNA-PK-Ku-eIF2-NF90-NF45 complex, the 10,000 random mean z score differences follow a normal distribution (mean = 0, s.d. = 0.228). Therefore, the observed mean z score difference of 0.98 between the N and LG groups corresponds to *p* = 0.002 (Fig. 6C).

Finally, applying the cutoffs of *p* < 0.05 and mean z score difference of > 0.5 in absolute value, 38 (34 up and 4 down) and 16 (15 up and 1 down) protein complexes were found to be differentially assembled in the LG and HG groups, respectively, compared with the N group (Fig. 6D and Table S9). Notably, nearly all of the differentially assembled protein complexes are not significantly differentially expressed (Table S9 versus Fig. 4B). After the removal of redundancy (*i.e.*, protein complexes comprising the same set of quantified proteins), 30 (27 up and 3 down) and 14 (13 up and 1 down) protein complexes were differentially assembled in the LG and HG groups, respectively (Table 1). Intriguingly, the protein complexes with increased assembly levels in the LG group (vs N) are mainly nuclear protein complexes, including those involved in chromatin remodeling, DNA damage response, as well as RNA synthesis and processing (Table 1 and Fig. S3). In comparison, the protein complexes with increased assembly levels in the HG group (vs N) are mainly subcomplexes of mitochondrial complex I (NADH-ubiquinone oxidoreductase) and Nucleosome Remodeling Deacetylase (NuRD) complex (Table 1).

**Table 1.**
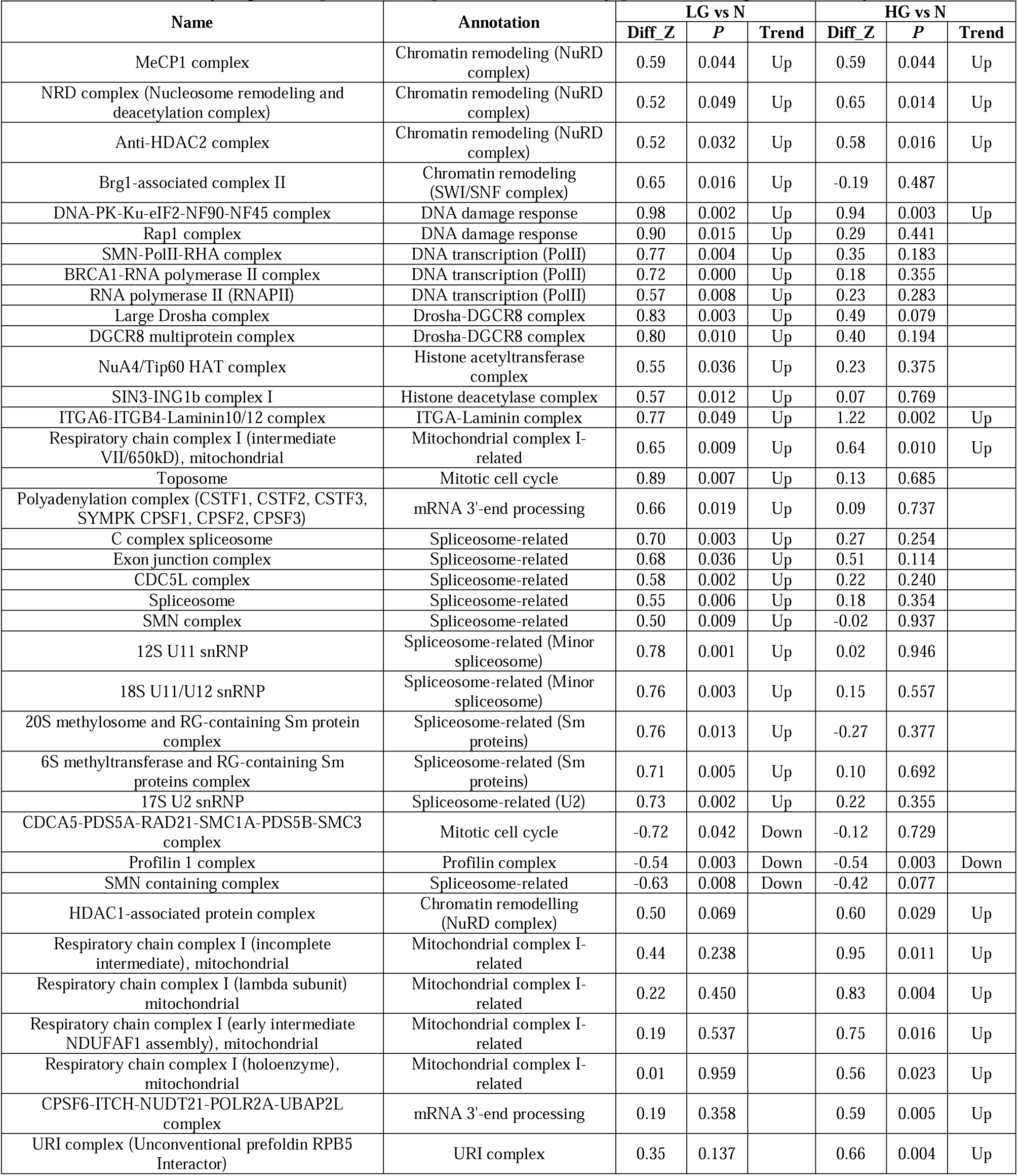
Differentially regulated protein complexes identified by protein co-regulation analysis

**Figure 4.**
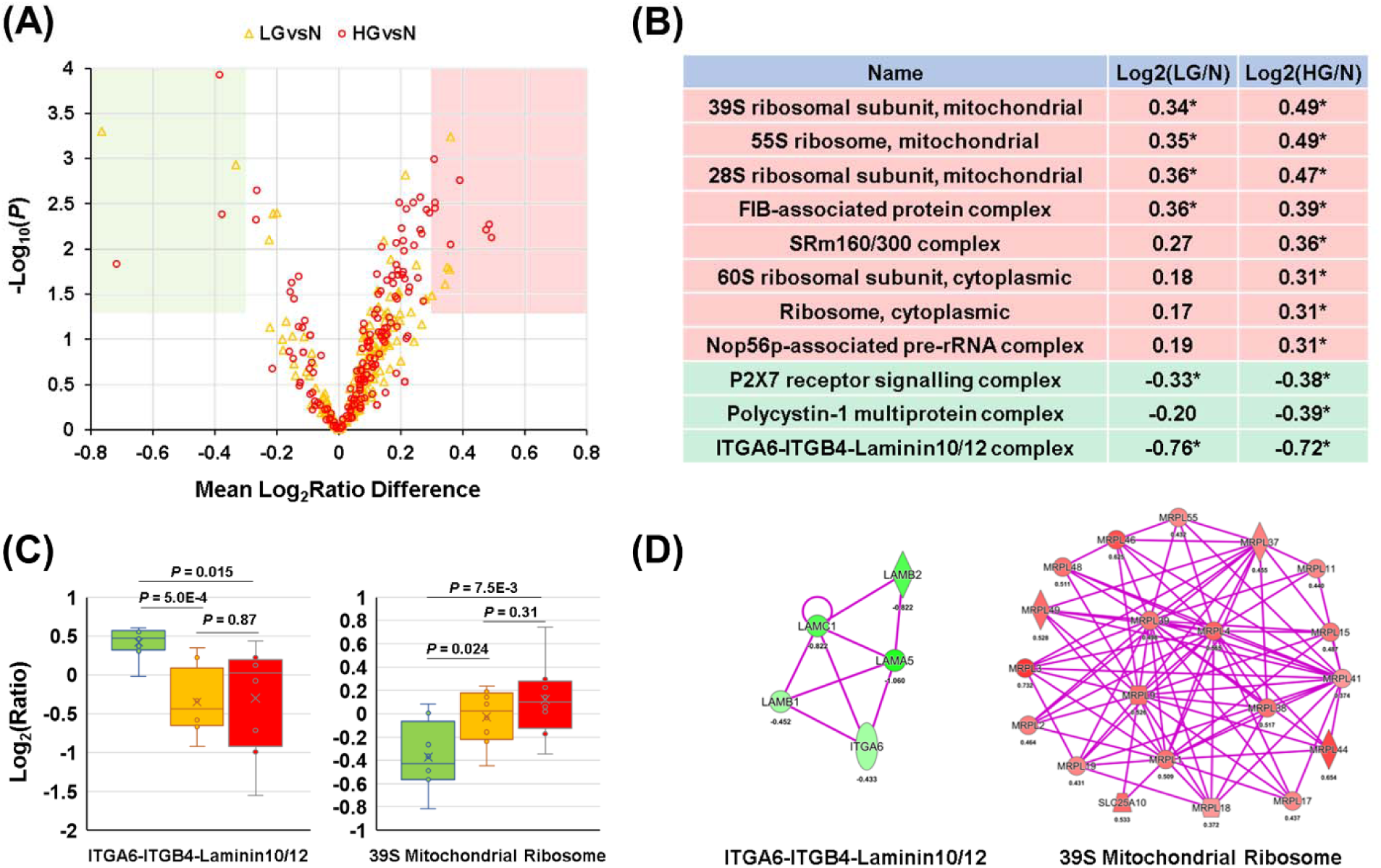
CORUM protein complexes differentially expressed in LG and HG PCa, compared with N samples. **(A)** Volcano plot showing log_2_-transformed fold changes plotted against negative log10-transformed *p* values for the 179 CORUM protein complexes. **(B)** List of protein complexes significantly (*p* < 0.05, indicated by *) up-or down-regulated in LG and/or HG PCa, compared with N samples. **(C)** Boxplots showing the differential expression of the ITGA6-ITGB4-Laminin10/12 complex and the 39S Mitochondrial Ribosomal Subunit, which are the most remarkably downregulated and upregulated in HG PCa (vs N), respectively. **(D)** Direct PPI networks of quantified proteins belonging to the ITGA6-ITGB4Laminin10/12 complex (left) and the 39S Mitochondrial Ribosomal Subunit (right). The numbers under gene names indicate the log_2_ (HG/N) ratios.

## Discussion

### Proteins differentially expressed in PCa versus normal prostate tissue

In this study, nearly 400 proteins were found to be differentially expressed between PCa and normal prostate tissue. In comparison, only 33 proteins were differentially expressed between low-grade and high-grade PCa specimens, even though the two groups have very different patient outcomes [41, 42]. Nevertheless, this is consistent with the finding of another global proteomic study of primary PCa tissue samples, which showed that only a small number of proteins were differentially expressed between low-risk and high-risk PCa groups [10]. It remains enigmatic how the modest proteomic changes lead to different PCa aggressiveness and distinct patient outcomes. However, it is possible that only a small subpopulation of PCa cells (*e.g.*, PCa stem cells) are responsible for PCa metastasis and drug resistance. These cells might have more profound proteomic changes than the bulk of PCa cells in high-grade PCa, in comparison to PCa cells in low-grade PCa.

Compared with normal prostate, the most remarkably up-and down-regulated proteins in low-grade PCa were identified as PYCR1 and MAMDC2, respectively. PYCR1 is a metabolic enzyme that catalyzes the NAD(P)H-dependent conversion of pyrroline-5-carboxylate to proline [46]. Previous studies showed that 1) compared with normal prostate, PYCR1 was significantly upregulated in PCa at both mRNA and protein levels, 2) the expression levels of PYCR1 were significantly associated with Gleason scores, and 3) PYCR1 is involved in PCa cell proliferation and colony formation [47, 48]. MAMDC2 is a poorly characterized proteoglycan containing four MAM domains, which are commonly found in surface receptors [49]. To our knowledge, no other studies reported that MAMDC2 was remarkably downregulated in low-grade PCa, compared with normal prostate.

Compared with normal prostate, the most dramatically up-and down-regulated proteins in high-grade PCa were identified as SPON2 and GSTM1, respectively. SPON2 is an extracellular matrix (ECM)protein belonging to the F-Spondin family. It was found to be a candidate serum and histological diagnostic biomarker for PCa and a candidate prognostic biomarker for colorectal cancer [50–53]. GSTM1 encodes a mu class cytoplasmic glutathione-*S*-transferase, which functions in cellular detoxification of many carcinogens. A recent meta-analysis suggested that GSTM1 deletion was significantly associated with PCa in overall, Asian, Eurasian, and American populations [54].

Compared with low-grade PCa, the most notably up-and down-regulated proteins in high-grade PCa were identified as PDSS2 and CD38, respectively. PDSS2, an enzyme that synthesizes the prenyl side-chain of coenzyme Q, was included in the Promark panel for the prediction of PCa aggressiveness and lethality [55, 56]. CD38, a cyclic ADP-ribose synthase, is the main NAD’ase in cells. Recent studies indicated that 1) decreased expression of CD38 in luminal progenitor cells can initiate PCa and is linked to lower overall survival, 2) CD38 expression inversely correlates with PCa progression, and 3) CD38 inhibits PCa proliferation by reducing cellular NAD^+^ pools [57, 58].

### The putative PPI networks of DEPs

The present study revealed that, compared with normal prostate, both low-grade and high-grade PCa have decreased expression of many ECM and plasma membrane proteins, which form large putative PPI networks. The ECM remodeling is probably due to increased proteolysis by matrix metalloproteinases and extracellular vesicles [59]. The most striking difference between low-grade and high-grade PCa is that only small PPI networks were more abundant in the former group, whereas a large PPI network was upregulated in the latter group. It suggests that high-grade PCa cells may potentially coordinate different protein complexes to achieve higher aggressiveness.

### Differentially expressed CORUM protein complexes

A prominent finding is the over-production of mitochondrial ribosomes in low-grade PCa and, to a greater extent, in high-grade PCa, compared with normal prostate. To our knowledge, the over-production of mitochondrial ribosomes in primary PCa has not yet been reported, although Iglesias-Gato *et al.* recently showed that mitochondrial protein content was upregulated in primary PCa and a subset of metastatic PCa [10, 14]. In human cells, mitochondrial ribosomes synthesize proteins encoded by 13 mitochondrial DNA-encoded genes. Recent studies suggested reduced mitochondrial DNA content in PCa [60, 61], so it is possible that PCa cells overexpress mitochondrial ribosomes to compensate the decrease of mitochondrial DNA content.

In addition, protein complexes related to ribosome biogenesis as well as cytoplasmic ribosomes were found to be overexpressed in high-grade PCa. This is consistent with previous findings that neoplastic prostate cells are characterized by enlarged, prominent nucleoli, and that nucleolar number and size are positively correlated with PCa malignancy [62].

### Differentially regulated protein complexes

The protein co-regulation analysis revealed that the assembly levels of the DNA-PK-Ku-eIF2-NF90NF45 complex and three subcomplexes of the chromatin remodeling NuRD complex were significantly increased in both low-grade and high-grade PCa, compared with normal prostate. Notably, these complexes play crucial roles in DNA damage repair [44, 45, 63]. The increased assembly levels of the complex may reflect higher genotoxic stress, which results in genome instability, in PCa cells than in normal prostate cells.

The co-regulation analysis also showed that the assembly levels of many nuclear complexes, especially those involved in RNA transcription and processing, are significantly increased in low-grade PCa but not in high-grade PCa, compared with normal prostate. It is unclear why their assembly levels in high-grade PCa did not reach the levels in low-grade PCa. One possibility is that high-grade PCa samples are more heterogenous and a subset is not dependent on increased assembly of these nuclear complexes.

The most prominent complexes with higher assembly levels in high-grade PCa, compared with normal prostate and low-grade PCa, are mitochondrial complex I and its subcomplexes (Table 1). As the largest complex of the mitochondrial electron transport chain, mitochondrial complex I contributes ∽40% of the proton motive force required for mitochondrial ATP synthesis [64]. Moreover, via modulating the NAD+/NADH ratio, mitochondrial complex I controls the synthesis of aspartate, a precursor of purine and pyrimidine synthesis. Although still controversial, epidemiological studies demonstrated that Metformin, a mitochondrial complex I inhibitor, reduces incidence and mortality of PCa patients [65]. It is possible that a combination of Metformin with androgen deprivation therapy may particularly benefit PCa patients with higher assembly levels of mitochondrial complex I in PCa cells.

### Limitations

Similar to all other comprehensive proteomic studies of prostate tissue specimens, the sample size in this study is relatively small. In addition, other categories of prostate tissue specimens, such as those of benign prostatic hyperplasia, PCa with Gleason scores of 7, and more importantly metastatic PCa, were not investigated. Nonetheless, larger-scale analysis of prostate tissue specimens in the near future will clarify the landscape of global protein complex changes during PCa development and progression, leading to the discovery of novel biomarkers and drug targets for precision management of PCa patients.

## Conclusions

In summary, TMT-SPS-MS3 profiling of clinical prostate tissue specimens, followed by differential expression and co-regulation analyses, led to the discovery of protein complexes deregulated in primary PCa. Compared with normal prostate, low-grade PCa tissue samples have 1) more abundant mitochondrial ribosomes and nuclear fibrillarin (FIB)-associated protein complex, 2) higher assembly levels of some nuclear protein complexes such as those involved in chromatin remodeling, DNA damage response, RNA synthesis, and RNA processing, 3) decreased abundance of protein subnetworks localized in extracellular space, plasma membrane, and cytoplasm, as well as of two CORUM complexes, and 4) decreased assembly levels of three CORUM complexes. In addition, compared with normal prostate, high-grade PCa tissue samples have 1) more abundant cytoplasmic and nuclear protein subnetworks as well as mitochondrial ribosomes, cytosolic ribosomes, ribosome biogenesis-related protein complexes, and an RNA splicing-related complex, 2) increased assembly of mitochondrial complex I and NuRD complexes, 3) decreased abundance of protein subnetworks localized in extracellular space, plasma membrane, and cytoplasm as well as of three CORUM complexes, and 4) decreased assembly of the profilin 1 complex. To our knowledge, this study represents the first comprehensive analysis of protein complexes in PCa tissue specimens. It is expected to enhance our understanding of the changes of protein complexes along PCa progression and lead to the identification of novel biomarkers and therapeutic targets for precision management of PCa patients.

## Declarations

### Ethics approval and consent to participate

The use of the prostate tissue specimens was approved by the Institutional Review Board of Cedars-Sinai Medical Center.

### Consent for publication

All authors consent to the publication of this manuscript.

### Availability of data and material

The mass spectrometry proteomics data have been deposited to the ProteomeXchange Consortium (http://proteomexchange.org) via the PRIDE partner repository [33] with database identifier PXD010744.

### Authors’ contributions

BZ and WY conceived and designed the study. BZ, YY, and YW carried out the experiments and contributed to the data acquisition. BZ performed database searching for protein identification and quantification. BZ, SY, and WY conducted statistical and bioinformatic analysis. BZ, MF, and WY interpreted results. BZ, SY, MF, and WY drafted the manuscript. All authors read and approved the final manuscript.

## Acknowledgements

We thank the Cedars-Sinai Biobank and Translational Pathology core for providing the OCT-embedded prostate tissue specimens.

## Abbreviations

DEP: Differentially expressed proteins
FASP: Filter-aided sample preparation
FDR: False discovery rate
FFPE: Formalin-fixed and paraffin-embedded
HG: High-grade
IPA: Ingenuity Pathway Analysis
LC: Liquid chromatography
LG: Low-grade
MALDI: Matrix-assisted laser desorption/ionization
MS: Mass spectrometry
N: Normal
OCT: Optimal cutting temperature
PCa: Prostate cancer
PPI: Protein-protein interaction
SPS: Synchronous precursor selection
2DE: Two-dimensional electrophoresis
TMT: tandem mass tag

## References

1. Torre LA, Bray F, Siegel RL, Ferlay J, Lortet-tieulent J, Jemal A. Global Cancer Statistics, 2012. CA a cancer J. Clin. 2015;65:87–108.

2. Siegel RL, Miller KD, Jemal A. Cancer statistics, 2018. CA. Cancer J. Clin. 2018;68:7–30.

3. Frank S, Nelson P, Vasioukhin V. Recent advances in prostate cancer research: large-scale genomic analyses reveal novel driver mutations and DNA repair defects. F1000Research. 2018;7:1173.

4. Roychowdhury S, Chinnaiyan AM. Translating cancer genomes and transcriptomes for precision oncology. CA. Cancer J. Clin. 2016;66:75–88.

5. You S, Knudsen BS, Erho N, Alshalalfa M, Takhar M, Al-deen Ashab H, et al. Integrated Classification of Prostate Cancer Reveals a Novel Luminal Subtype with Poor Outcome. Cancer Res. 2016;76:4948–58.

6. Latonen L, Afyounian E, Jylhä A, Nättinen J, Aapola U, Annala M, et al. Integrative proteomics in prostate cancer uncovers robustness against genomic and transcriptomic aberrations during disease progression. Nat. Commun. Springer US; 2018;9:1176.

7. Zhang B, Wang J, Wang X, Zhu J, Liu Q, Shi Z, et al. Proteogenomic characterization of human colon and rectal cancer. Nature. 2014;513:382–7.

8. Murray HC, Dun MD, Verrills NM. Harnessing the power of proteomics for identification of oncogenic, druggable signalling pathways in cancer. Expert Opin. Drug Discov. 2017;12:431–47.

9. Mantsiou A, Vlahou A, Zoidakis J. Tissue proteomics studies in the investigation of prostate cancer. Expert Rev. Proteomics. Taylor & Francis; 2018;00:1–19.

10. Iglesias-Gato D, Wikström P, Tyanova S, Lavallee C, Thysell E, Carlsson J, et al. The Proteome of Primary Prostate Cancer. Eur. Urol. 2016;69:942–52.

11. Staunton L, Tonry C, Lis R, Espina V, Liotta L, Inziatari R, et al. Pathology-driven Comprehensive Proteomic Profiling of the Prostate Cancer Tumour Microenvironment. Mol. Cancer Res. 2017;molcanres.0358.2016.

12. Müller A-K, Föll M, Heckelmann B, Kiefer S, Werner M, Schilling O, et al. Proteomic Characterization of Prostate Cancer to Distinguish Nonmetastasizing and Metastasizing Primary Tumors and Lymph Node Metastases. Neoplasia. The Authors; 2018;20:140–51.

13. Guo T, Li L, Zhong Q, Rupp NJ, Charmpi K, Wong CE, et al. Multi-region proteome analysis quantifies spatial heterogeneity of prostate tissue biomarkers. Life Sci. Alliance. 2018;1:e201800042.

14. Iglesias-Gato D, Thysell E, Tyanova S, Crnalic S, Santos A, Lima TS, et al. The proteome of prostate cancer bone metastasis reveals heterogeneity with prognostic implications. Clin. Cancer Res. 2018;clincanres.1229.2018.

15. Alberts B. The Cell as a Collection Overview of Protein Machines: Preparing theNext Generation of Molecular Biologists. Cell. 1998;92:1–4.

16. Havugimana PC, Hu P, Emili A. Protein complexes, big data, machine learning and integrative proteomics: lessons learned over a decade of systematic analysis of protein interaction networks. Expert Rev. Proteomics. Taylor & Francis; 2017;14:845–55.

17. Wang PI, Marcotte EM. It’s the machine that matters: Predicting gene function and phenotype from protein networks. J. Proteomics. Elsevier B.V.; 2010;73:2277–89.

18. Ankney JA, Muneer A, Chen X. Relative and Absolute Quantitation in Mass Spectrometry–Based Proteomics. Annu. Rev. Anal. Chem. 2018;11:49–77.

19. Thompson A, Schäfer J, Kuhn K, Kienle S, Schwarz J, Schmidt G, et al. Tandem mass tags: a novel quantification strategy for comparative analysis of complex protein mixtures by MS/MS. Anal. Chem. 2003;75:1895–904.

20. McAlister GC, Huttlin EL, Haas W, Ting L, Jedrychowski MP, Rogers JC, et al. Increasing the multiplexing capacity of TMTs using reporter ion isotopologues with isobaric masses. Anal. Chem. 2012;84:7469–78.

21. Ting L, Rad R, Gygi SP, Haas W. MS3 eliminates ratio distortion in isobaric multiplexed quantitative proteomics. Nat. Methods. United States; 2011;8:937–40.

22. McAlister GC, Nusinow DP, Jedrychowski MP, W?hr M, Huttlin EL, Erickson BK, et al. MultiNotch MS3 Enables Accurate, Sensitive, and Multiplexed Detection of Differential Expression across Cancer Cell Line Proteomes. Anal. Chem. United States; 2014;86:7150–8.

23. O’Connell JD, Paulo JA, O’Brien JJ, Gygi SP. Proteome-Wide Evaluation of Two Common Protein Quantification Methods. J. Proteome Res. 2018;17:1934–42.

24. Lapek JD, Greninger P, Morris R, Amzallag A, Pruteanu-Malinici I, Benes CH, et al. Detection of dysregulated protein-association networks by high-throughput proteomics predicts cancer vulnerabilities. Nat. Biotechnol. Nature Publishing Group; 2017;35:983–9.

25. Holfeld A, Valdés A, Malmström PU, Segersten U, Lind SB. Parallel Proteomic Workflow for Mass Spectrometric Analysis of Tissue Samples Preserved by Different Methods. Anal. Chem. 2018;90:5841–9.

26. P.D. P, V.A. P, R.L. S, M.A. G, K.K. W, T.L. F, et al. Residual tissue repositories as a resource for population-based cancer proteomic studies. Clin. Proteomics. BioMed Central; 2018;15:26.

27. Zhang W, Sakashita S, Taylor P, Tsao MS, Moran MF. Comprehensive proteome analysis of fresh frozen and optimal cutting temperature (OCT) embedded primary non-small cell lung carcinoma by LC-MS/MS. Methods. Elsevier Inc.; 2015;81:50–5.

28. Qu Y, Zhou B, Yang W, Han B, Yu-Rice Y, Gao B, et al. Transcriptome and proteome characterization of surface ectoderm cells differentiated from human iPSCs. Sci. Rep. Nature Publishing Group; 2016;6:32007.

29. Gonsky R, Fleshner P, Deem RL, Biener-Ramanujan E, Li D, Potdar AA, et al. Association of Ribonuclease T2 Gene Polymorphisms With Decreased Expression and Clinical Characteristics of Severity in Crohn’s Disease. Gastroenterology. 2017;153:219–32.

30. Wang Y, Yang F, Gritsenko MA, Wang Y, Clauss T, Liu T, et al. Reversed-phase chromatography with multiple fraction concatenation strategy for proteome profiling of human MCF10A cells. Proteomics. 2011;11:2019–26.

31. Cox J, Mann M. MaxQuant enables high peptide identification rates, individualized p.p.b.-range mass accuracies and proteome-wide protein quantification. Nat. Biotechnol. 2008;26:1367–72.

32. Cox J, Neuhauser N, Michalski A, Scheltema RA, Olsen J V, Mann M. Andromeda: a peptide search engine integrated into the MaxQuant environment. J. Proteome Res. United States; 2011;10:1794–805.

33. Vizcaíno JA, Csordas A, Del-Toro N, Dianes JA, Griss J, Lavidas I, et al. 2016 update of the PRIDE database and its related tools. Nucleic Acids Res. 2016;44:D447–56.

34. Tyanova S, Temu T, Sinitcyn P, Carlson A, Hein MY, Geiger T, et al. The Perseus computational platform for comprehensive analysis of (prote)omics data. Nat. Methods. 2016;

35. Erickson BK, Rose CM, Braun CR, Erickson AR, Knott J, McAlister GC, et al. A Strategy to Combine Sample Multiplexing with Targeted Proteomics Assays for High-Throughput Protein Signature Characterization. Mol. Cell. 2017;65:361–70.

36. Storey JD. A direct approach to false discovery rates. J. R. Stat. Soc. Ser. B (Statistical Methodol. 2002;64:479–98.

37. Huang DW, Sherman BT, Lempicki RA. Systematic and integrative analysis of large gene lists using DAVID bioinformatics resources. Nat. Protoc. 2009;4:44–57.

38. Ryan CJ, Kennedy S, Bajrami I, Matallanas D, Lord CJ. A Compendium of Co-regulated Protein Complexes in Breast Cancer Reveals Collateral Loss Events. Cell Syst. Elsevier Inc.; 2017;5:399–409.e5.

39. Fukushima A. DiffCorr: An R package to analyze and visualize differential correlations in biological networks. Gene. Elsevier B.V.; 2013;518:209–14.

40. McKenzie AT, Katsyv I, Song WM, Wang M, Zhang B. DGCA: A comprehensive R package for Differential Gene Correlation Analysis. BMC Syst. Biol. BMC Systems Biology; 2016;10:1–25.

41. Matoso A, Epstein JI. Grading of Prostate Cancer: Past, Present, and Future. Curr. Urol. Rep. 2016;17:25.

42. Eggener SE, Scardino PT, Walsh PC, Han M, Partin AW, Trock BJ, et al. Predicting 15-year prostate cancer specific mortality after radical prostatectomy. J. Urol. American Urological Association Education and Research, Inc.; 2011;185:869–75.

43. Ruepp A, Waegele B, Lechner M, Brauner B, Dunger-Kaltenbach I, Fobo G, et al. CORUM: the comprehensive resource of mammalian protein complexes--2009. Nucleic Acids Res. 2010;38:D497–501.

44. Ting NS, Kao PN, Chan DW, Lintott LG, Lees-Miller SP. DNA-dependent protein kinase interacts with antigen receptor response element binding proteins NF90 and NF45. J. Biol. Chem. 1998;273:2136–45.

45. Fell VL, Schild-Poulter C. The Ku heterodimer: Function in DNA repair and beyond. Mutat. Res. Rev. Mutat. Res. Elsevier B.V.; 2015;763:15–29.

46. Yeh GC, Harris SC, Phang JM. Pyrroline-5-carboxylate reductase in human erythrocytes. J. Clin. Invest. 1981;67:1042–6.

47. Ernst T, Hergenhahn M, Kenzelmann M, Cohen CD, Bonrouhi M, Weninger A, et al. Decrease and gain of gene expression are equally discriminatory markers for prostate carcinoma: A gene expression analysis on total and microdissected prostate tissue. Am. J. Pathol. 2002;160:2169–80.

48. Zeng T, Zhu L, Liao M, Zhuo W, Yang S, Wu W, et al. Knockdown of PYCR1 inhibits cell proliferation and colony formation via cell cycle arrest and apoptosis in prostate cancer. Med. Oncol. Springer US; 2017;34:1–9.

49. Beckmann G, Bork P. An adhesive domain detected in functionally diverse receptors. Trends Biochem. Sci. 1993;18:40–1.

50. Qian X, Li C, Pang B, Xue M, Wang J, Zhou J. Spondin-2 (SPON2), a More Prostate-Cancer-Specific Diagnostic Biomarker. Addison CL, editor. PLoS One. 2012;7:e37225.

51. Lucarelli G, Rutigliano M, Bettocchi C, Palazzo S, Vavallo A, Galleggiante V, et al. Spondin-2, a secreted extracellular matrix protein, is a novel diagnostic biomarker for prostate cancer. J. Urol. Elsevier Ltd; 2013;190:2271–7.

52. Zhu B-P, Guo Z-Q, Lin L, Liu Q. Serum BSP, PSADT, and Spondin-2 levels in prostate cancer and the diagnostic significance of their ROC curves in bone metastasis. Eur. Rev. Med. Pharmacol. Sci. 2017;21:61–7.

53. Schmid F, Wang Q, Huska MR, Andrade-Navarro MA, Lemm M, Fichtner I, et al. SPON2, a newly identified target gene of MACC1, drives colorectal cancer metastasis in mice and is prognostic for colorectal cancer patient survival. Oncogene. Nature Publishing Group; 2016;35:5942–52.

54. Malik SS, Kazmi Z, Fatima I, Shabbir R, Perveen S, Masood N. Genetic Polymorphism of GSTM1 and GSTT1 and Risk of Prostatic Carcinoma - a Meta-analysis of 7,281 Prostate Cancer Cases and 9,082 Healthy Controls. Asian Pac. J. Cancer Prev. 2016;17:2629–35.

55. Margariti A, Winkler B, Karamariti E, Zampetaki A, Tsai T -n., Baban D, et al. Direct reprogramming of fibroblasts into endothelial cells capable of angiogenesis and reendothelialization in tissue-engineered vessels. Proc. Natl. Acad. Sci. 2012;109:13793–8.

56. Blume-Jensen P, Berman DM, Rimm DL, Shipitsin M, Putzi M, Nifong TP, et al. Biology of Human Tumors Development and clinical validation of an in situ biopsy-based multimarker assay for risk stratification in prostate cancer. Clin. Cancer Res. 2015;21:2591–600.

57. Liu X, Grogan TR, Hieronymus H, Hashimoto T, Mottahedeh J, Cheng D, et al. Low CD38 Identifies Progenitor-like Inflammation-Associated Luminal Cells that Can Initiate Human Prostate Cancer and Predict Poor Outcome. Cell Rep. ElsevierCompany.; 2016;17:2596–606.

58. Chmielewski JP, Bowlby SC, Wheeler FB, Shi L, Sui G, Davis AL, et al. CD38 Inhibits Prostate Cancer Metabolism and Proliferation by Reducing Cellular NAD+ Pools. Mol. Cancer Res. 2018;molcanres.0526.2017.

59. Karamanos NK, Theocharis AD, Neill T, Iozzo R V. Matrix modeling and remodeling: A biological interplay regulating tissue homeostasis and diseases. Matrix Biol. International Society of Matrix Biology; 2018;

60. Koochekpour S, Marlowe T, Singh KK, Attwood K, Chandra D. Reduced Mitochondrial DNA Content Associates with Poor Prognosis of Prostate Cancer in African American Men. PLoS One. 2013;8.

61. Kalsbeek AMF, Chan EKF, Grogan J, Petersen DC, Jaratlerdsiri W, Gupta R, et al. Altered mitochondrial genome content signals worse pathology and prognosis in prostate cancer. Prostate. 2018;78:25–31.

62. Wilkinson N, Buckley CH, Chawner L, Fox H. Nucleolar organiser regions in normal, hyperplastic, and neoplastic endometria. Int. J. Gynecol. Pathol. 1990;9:55–9.

63. Basta J, Rauchman M. The Nucleosome Remodeling and Deacetylase Complex in Development and Disease. Transl. Epigenetics to Clin. Elsevier Inc.; 2017.

64. Urra FA, Muñoz F, Lovy A, Cárdenas C. The Mitochondrial Complex(I)ty of Cancer. Front. Oncol. 2017;7:1–8.

65. Whitburn J, Edwards CM, Sooriakumaran P. Metformin and Prostate Cancer: a New Role for an Old Drug. Curr. Urol. Rep. Current Urology Reports; 2017;18:1–7.

